# Connecting human voice profiling to genomics: A predictive algorithm for linking speech phenotypes to genetic microdeletion syndromes

**DOI:** 10.1101/2022.05.23.493126

**Authors:** Rita Singh

## Abstract

Changes in vocal acoustic patterns are known to correlate with the occurrence of several diseases and syndromes, many of which do not directly affect the structures or processes that control voice production. In such cases, it is difficult to support the existence of correlated changes in voice. This paper presents a methodology for identifying potential genomic bases for such correlations, by finding links between specific genes involved in the conditions under study, and those involved in voice, speech or language generation. Syndromes associated with chromosomal microdeletions are examined as an illustrative case, with focus on their linkage to the FOXP2 gene which has been strongly implicated in speech and language disorders. A novel path-finding graph algorithm to detect pathway chains that connect the the former to the latter is proposed. Statistical analysis of ensembles of “voice” chains detected by this algorithm indicates that they are predictive of speech phenotypes for the syndromes. Algorithmic findings are validated against clinical findings in the literature pertaining to the actual speech phenotypes that have been found to be associated with these syndromes. This methodology may also potentially be used to predict the existence of voice biomarkers in naїve cases where the existence of voice biomarkers has not already been established.

## I. Introduction

Aside from diseases that affect the biological structures and processes involved in voice production, myriad other factors are known to influence voice. The ensuing changes in the voice signal are the biomarkers that give us information about the causal factors, and allow us to infer their nature through voice analysis. Such relationships form the basis for Artificial Intelligence (AI) based voice profiling techniques that attempt to deduce a speaker’s bio-relevant and environmental parameters from voice. However, virtually all research on voice profiling, diagnostics and biometrics is predicated on clinically observed or statistically inferred relationships between changes in voice, and the corresponding factors that are thought to be causal. The relationships that are chanced upon in this manner provide the basis for building predictive AI (machine learning or rule-based) mechanisms that can deduce the underlying factors (that potentially influence voice) through voice analysis.

For example, it is known that smoking affects voice. To establish this, a *human-observation-based approach* would be: a) an audiological one based on hearing the voices of smokers to determine if they deviate from those of non-smokers in an acoustic sense, and/or b) a visual one where an analyst studies the spectrogram (or some other visual representation) of the speech signal to find patterns that distinguish one class of recordings from another. The spectrogram in this case is a “feature representation.” A *statistical approach*, on the other hand, would gather examples of speech recordings from people who smoke and those who don’t, extract feature representations from the recordings, and find significant differences in the statistics of these features obtained from the two sets of recordings. Alternatively, a classifier model may be trained to discriminate between voice samples from smokers and non-smokers. If high test accuracies are achieved in this task, the existence of a biomarker for smoking in voice is indicated. This is a purely data-driven approach for establishing the existence of biomarkers in voice.

The problem with these approaches is that neither is scalable. The number of factors that can influence the human persona is virtually infinite. Human observations are limited to the effects that are perceptually discernible in voice, and data-based discovery is confined and limited by the availability of representative data. This paper provides a more formal methodology for establishing the existence of biomarkers, and for identifying which factors are likely to affect voice and which are not. The methodology is based on genomic considerations, explained below.

### I-A. A genomic-based approach to detect the existence of biomarkers

The working hypothesis for this paper, and one that has also recently been proposed in the context of voice profiling [1], is that if a given factor exerts an influence on the speaker, and if pathways of biological effects can be traced from that influence to the speaker’s voice production system, then voice *must* be affected (and must carry biomarkers for the factor). The methodology proposed herein is a literal test of this hypothesis, in that it traces biological pathways from cause to effect, to establish the existence of biomarkers.

For this, we begin with the genetic underpinnings of human vocal capabilities. In this context, it is important to differentiate between voice production and speech production. The former refers to the production of acoustic energy in any form within the vocal tract, and the latter refers to the modulation of the acoustic signals thus produced, to form words and sentences in a language used for interpersonal communication.

The question of whether human voice and speech production capabilities have a genetic basis has been widely explored. In the quest for identifying and delineating the genes responsible for functional speech in humans, recent studies have highlighted the importance of FOXP2, a protein-coding gene, that is thought to be involved in a variety of biological pathways and cascades that may regulate language development. It is autosomal dominant, and mutations in it cause speech and language disorders (OMIM: SPCH1). In this paper, we use FOXP2 as an exemplary case, noting that the methodology presented in its context is itself a generic one, and can be applied to any set of genes whose functions are relevant to the analysis at hand. For illustrative purposes, we proceed with the broad and simplifying assumption that any influence on speech and language is ultimately the phenotypic expression of FOXP2. The objective is now to:

- Formalize the methodology to find a link between an influencing factor, and this gene
- Validate the methodology
- Demonstrate its predictive potential

We accomplish these goals by reconciling the genomic findings for specific syndromes resultant from genetic aberrations, to clinical observations of the speech phenotype in each case. We choose to work with chromosomal microdeletions, and suggest a method to link the genes involved in the cytogenetic locations of the microdeletions to FOXP2, as a specific example. The connections are derived using a path-search algorithm applied to a graph composed from known biological pathways and the linkages between them.

### I-B. Anomalies in speech production

From a bio-mechanical perspective, Human speech is the result of two complex processes that happen simultaneously: one that produces sound – the pressure wave that we sense as the voice signal – and another that modulates this signal (through articulator movements) to produce speech – altering the voice signal’s frequency characteristics, and shaping it into sounds with unique identities that are uttered sequentially to form words and sentences in a language. The overall process of voice and speech production is driven and controlled by neuromuscular and cognitive factors to different degrees. It is also moderated to different degrees by feedback obtained through auditory pathways. Generally, diseases that affect these functions naturally also influence speech, and alter the characteristics of the voice signal to proportional degrees. In most cases, when reporting such changes, references to “speech” implicitly include voice, as we see below.

Changes in speech are categorically described in terms of six major aspects of speech production: respiration, phonation, articulation, resonance, evolution and prosody. In addition, terminology that relates to *voice quality* is often used to describe speech. Voice quality is however a subjective term, and comprises many constituents, or sub-qualities (e.g. nasal, breathy, rough, twangy etc.) that refer to the perceptual flavor of speech (or how a speaker’s voice sounds to the listener). Physical anomalies that affect the shape and tissue structure of the vocal tract cause changes in all of these aspects. Speech delays and language difficulties result from cognitive and learning disabilities. These and other intellectual disabilities affect articulation, evolution and prosody. Their effect on voice also manifests as changes in voice quality. Craniofacial anomalies affect the physical dimensions of the vocal tract structures, often restricting the movement of the articulators as a result, and causing speaking impairments. Motor problems affect articulation, phonation and respiration. These cause speech aberrations and also affect voice quality. Hearing problems disturb the feedback mechanisms involved in controlling speech production, and often lead to difficulties in prosody and articulation of speech.

In this paper, we do not focus on voice acoustic or quality characteristics separately. Instead, we focus exclusively on problems with speech (subsuming voice to some extent) as described under the OMIM (referring to the catalog Online Mendelian Inheritance in Man: https://www.omim.org/) category “Speech and Language disorders (SPCH1).” This category, however, is too broad and encompasses a wide range of speech problems (or speech phenotypes), such as delay in acquisition of speech abilities, retardation in speech development with age, speech anomalies resulting from language delay, expression, and articulation. For the purpose of this paper, it is necessary to make finer distinctions between speech phenotypes. The problem in doing so is that the language used in the literature to describe voice and speech related problems in the context of genetic syndromes is not standardized. For example, the terms “speech disorder”, “speech disturbance”, “speech anomalies”, “speech aberrations”, and “speech impairment” may each refer to a range of symptoms that may be overlapping to various degrees. For the purpose of this paper, it is therefore useful to map the broad range of speech phenotypes into the following categories that are sufficiently discriminatory in terms of the different aspects of speech production mentioned above, while still being limited in number:

#### 1) Absence of speech

Phrases (in clinical/scientific literature) referring to a) no development of speech capabilities, b) no expressive speech – mostly limited to vocalizations, c) almost absent speech, with severely limited vocabulary (0-4 words).

#### 2) Apraxia

Phrases referring to difficulty using language correctly while speaking, leading to speaking and communication difficulties.

#### 3) Delayed speech

Phrases referring to developmental delay, or the retarded development of the ability to speak, retarded acquisition of language skills and communication skills (ability to use a vocabulary correctly to communicate in a cogent manner)

#### 4) Dysarthric speech or Dysarthria

Phrases referring to speaking problems resultant from damaged, paralyzed, or weakened muscles of the articulators, caused by motor problems. Dysarthria results in slurred words, poor phonation etc. The speaker uses vocabulary as in normal speech, but finds it difficult to move the articulators (tongue, lips, jaw etc) correctly to form the proper sounds to utter it.

#### 5) Idiosyncratic speech

Phrases referring to poor conformance to cogent language, or incoherent language with articulation abnormalities.

#### 6) Impaired speech

Phrases referring to poor articulation, phonation, and difficulties that result in sparse and disfluent speech.

## II. Methodology

We study the syndromes resulting from chromosomal microdeletions, focusing on the phenotypic expressions of speech abilities. Chromosomal microdeletions are structural anomalies of chromosomes in which small sections of a chromosome are deleted or missing. The loss of the specific set of genes from the deleted section often results in phenotypic changes. An *implicated gene* is a gene in the deleted region of a chromosome, that is known to correlate well to the phenotypic expression of the syndrome represented by the deletion, as supported by microarray and other studies. Although the human genome is large in comparison to a typical chromosomal microdeletion region, only a specific set of deletions are compatible with life, or fetal survival. This set is small, and continues to expand with the addition of newly discovered deletions in surviving individuals who have the means to reach genetic testing facilities. Most known deletions are thus well documented in the literature, both from the genetic and medical perspectives. The genes associated with them are largely identified through microarray techniques. The choice of microdeletion syndromes for this study is driven by the fact that deletions represent simple but selective elimination of genes from chromosomes, which are expected to be easier to interpret for their phenotypic effect. They allow us to explore a binary presence/absence causation of phenotypes with respect to genes.

The methodology proposed herein analyzes ensembles of biological pathway *chains*, each of which connects a specific gene in the cytogenetic region of chromosomal microdeletion to the FOXP2 gene. A “biological pathway” here is as defined in standard terminology, referring to a physiological process at the cellular level that is enabled by the action of multiple genes that perform specific functions within the process.

We define a *pathway chain* as the sequential linkage of pathways, where links between pathways are shared genes (implicitly meaning that the molecules resultant from genes are shared – we will use the term “gene” with this implicit meaning in the context of pathways for brevity, going forward). For example,a pathway that signals for a cell to stop dividing when an injury to the nuclear DNA strand is being repaired may involve the co-ordinated chemical action of molecules that are formed by the transcription of multiple genes which perform different functions, and would also be connected to a repair pathway by necessity. Thus the two pathways can be considered to be links in a single pathway chain – they must share some genes that perform the function of relaying messages from one pathway to another. Such genes may also perform other functions that are essential to both pathways.

The assumption we make here is that if, for a gene, a chain from its pathway(s) (i.e., the biological pathways it contributes to) extends to pathways that influence voice production, then the phenotype resultant from the absence or aberrant functioning of the gene can be expected to include anomalies in speech production and voice characteristics.

### II-A. Voice chains

Our definition of a *voice chain* extends our definition of a pathway chain, in that the head of the chain must now necessarily be a pathway that includes a gene that influences voice or speech production, while the termination of the chain is not necessarily a biological pathway, but could include any given set of genes with a common characterization (such as a common cytogenetic location or function).

In this paper the voice-related gene chosen is FOXP2, but in other analyses, voice chains could involve other genes (e.g. as in [2]) without loss of generality. The terminal link in the chain is taken to be a genetic microdeletion syndrome. A voice chain thus establishes a relationship between an influencing factor – a genetic microdeletion syndrome in this case – and a corresponding effect on voice/speech production. We refer to a voice chain that includes a sequence of *α* pathways from the microdeletion region to the FOXP2 gene as a *level-α* voice chain (Figure 1). The specific genes on the microdeletion that link it to the voice chain are referred to as “chainlink” genes.

**Fig. 1:**
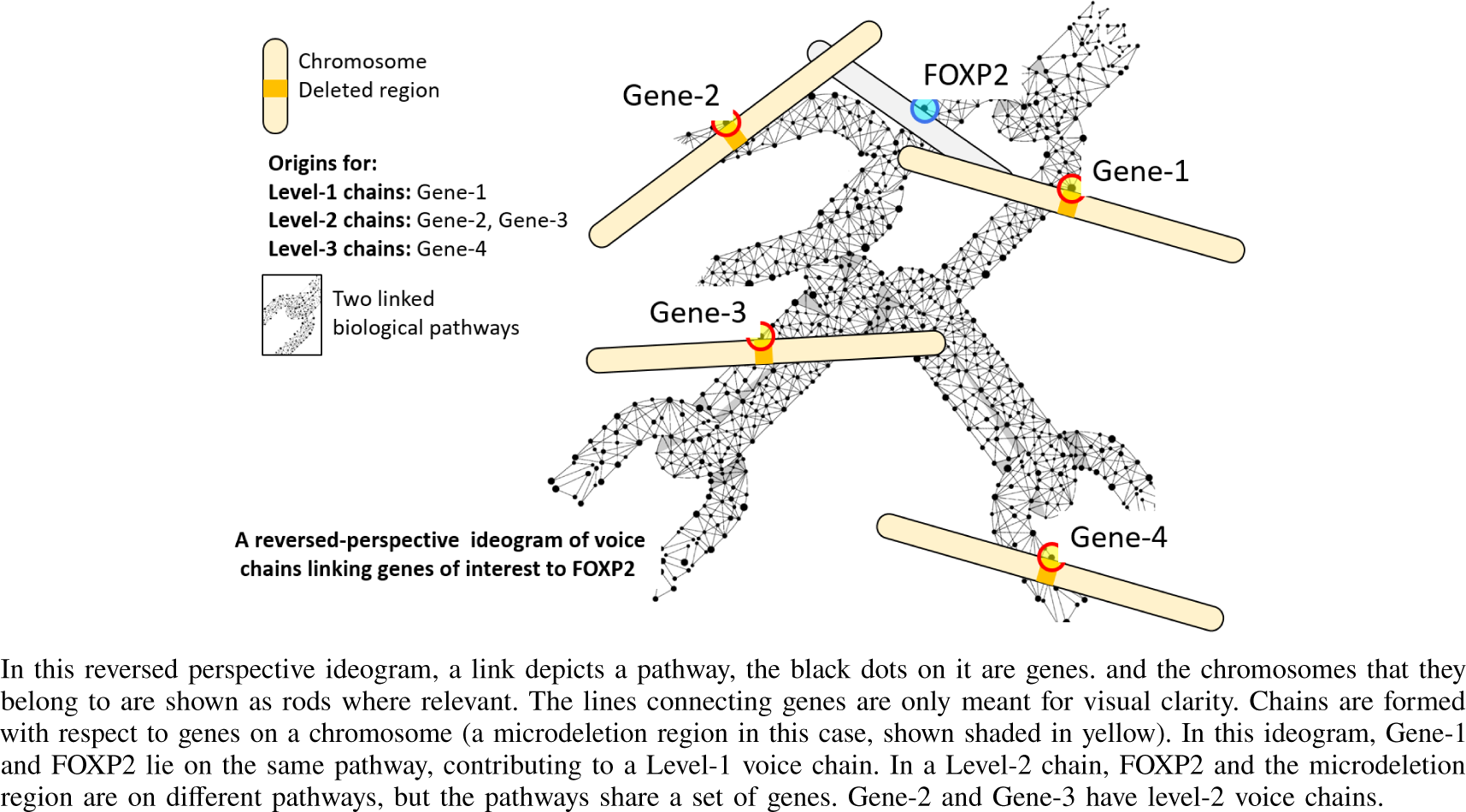
Voice chains of different levels.

In order to trace the genetic links between a microdeletion syndrome and voice, we first attempt to identify voice chains of different lengths that link to the genes in the microdeletion region. For this, we must find voice chains that link the FOXP2 gene to the syndrome, and identify the specific genes from the syndrome through which they are linked. We do so using the graph-search algorithm described below.

From our perspective, a biological pathway is ℬ represented by the set of genes it involves: ℬ := {*g* : *g* is a gene in the specified pathway }. Two pathways ℬ_1_ and ℬ_2_ are *linked* if there are genes that are common to both pathways, *i*.*e*. ℬ_1_ ∩ ℬ_2_ ≠ 0 Thus, the set of all pathways can be represented as a graph where the nodes are biological pathways, and two nodes are linked only if the corresponding pathways have common genes as illustrated in Figure 2a.

**Fig. 2:**
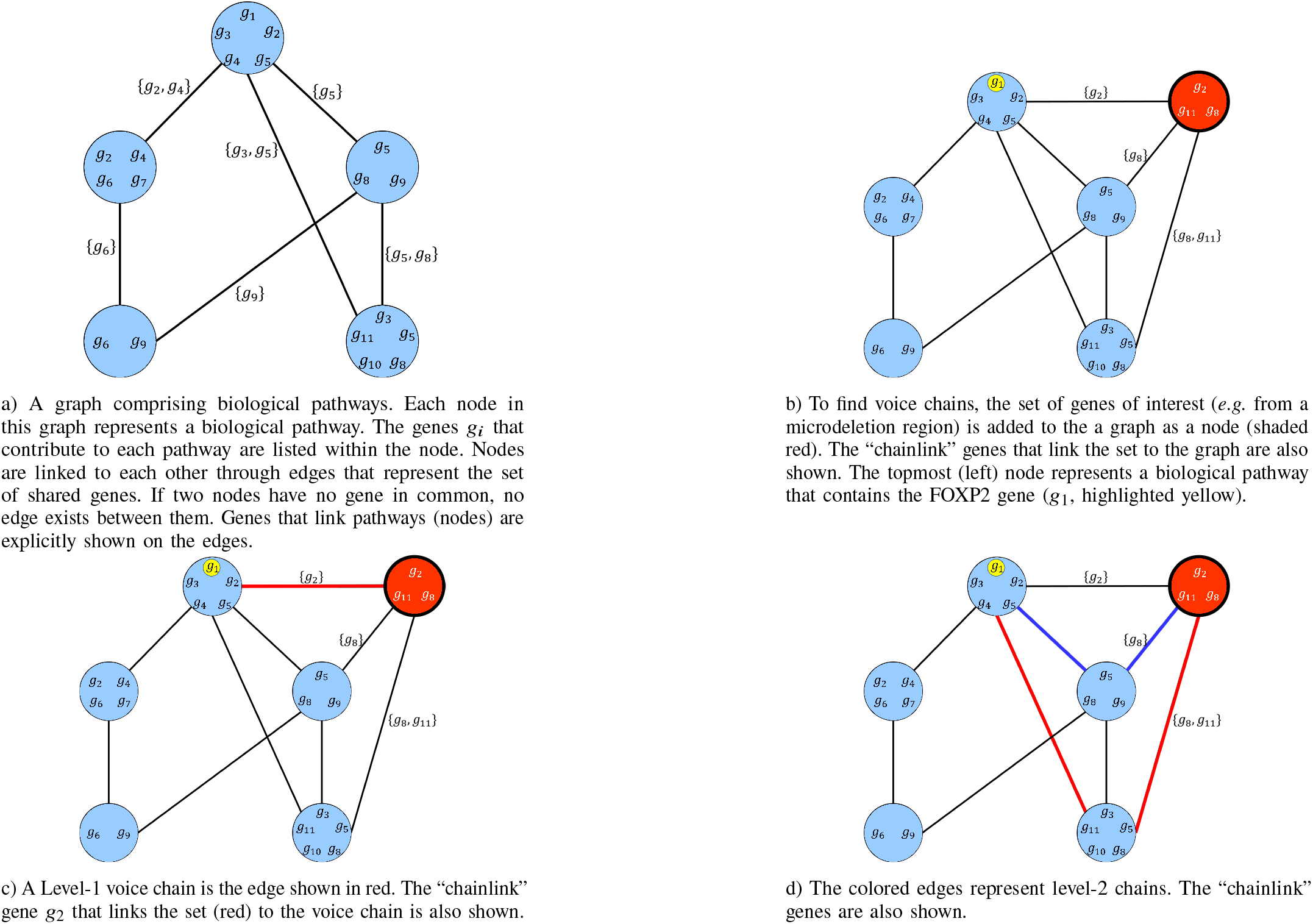
Voice chains of different types.

A pathway *chain* is any non-repeating sequence of pathways ℬ_1_ ℬ_2_ ℬ_3_ ℬ_*N*_ such that ℬ_i_∩ ℬ_i_+_1_ ≠ ∅, and ℬ_i_ ≠ ℬ_*j*_ for _i_ ≠ _*j*_, *i*.*e*. where every pair of adjacent pathways has common genes, and there are no closed loops in the chain. In terms of the graph (See Figure 2a), a pathway chain is any path between any two nodes in the graph. A *voice chain* 𝒱 is any chain 𝒱 = ℬ_*v*_ ℬ_2_ℬ_3_ … ℬ_*N*_ 𝒮 where the *head* node ℬ_*v*_ (and the head node alone) is a pathway that includes the FOXP2 gene, *i*.*e. FO X P*2 ∈ ℬ_*v*_ and the terminal node 𝒮 is a set of genes with common characterization, as mentioned earlier. The *length* of the chain, | 𝒱 | is the number of nodes *α* in the chain, not counting the terminal node 𝒮. We represent the set of voice chains of length *α* as 𝒱_*N α*_ (with the subscript _*N*_ indicating that it is a set and not a single chain). 𝒱_*N α*_ is thus the set of “level-*α*” chains. For the purpose of this paper, we will assume 𝒮 to be set of genes in a microdeletion region associated with a syndrome. Thus 𝒮 = {*g* : *g* is a gene in the microdeletion region}.

To find voice chains of the form ℬ_*v*_, ℬ_1_, … 𝒮 arising from the microdeletion region 𝒮 (which we will refer to as “syndrome” for brevity) we introduce it in the pathw y g (Figure 2b). Voice chains are now the paths from ℬ_*v*_ to 𝒮 (Figures 2c, 2d). A breadth-first algorithm described in Algorithm 1 is used to extract both, the voice chains and the genes through which the chains link to the syndrome. The outcome of the algorithm is the set of level-*α* voice chains 𝒱_*N α*_ for 1 ≤ *α* ≤ 2, as well as the sets 𝒢 [𝒮][*α*] comprising the sets of chainlink genes from each syndrome 𝒮 that links to voice chains of lengths *α*. We restrict ourselves to chains of length up to 2, since at greater lengths the chained influences cannot be disambiguated as indicated by prior studies in the (highly correlated) context of protein-protein interactomes, *e*.*g*. [3].

### II-B. Ensemble analysis

In the methodology we propose, for any microdeletion region 𝒮, we derive the set of chainlink genes within it for which level-*α* chains exist. The size and composition of this set can then be used in conjunction with the level of the voice chain to indicate the effect on voice phenotype (in later analysis). In general, we can work with any level-*α* voice chains in such an analysis; however we restrict ourselves to *α* = 1 and *α* = 2.

## III. Analysis

𝒱_*N α*_, *α* = 1, 2 were computed for a total of 82 microdeletion syndromes of chromosomes 1-20/22/X,Y. Genomic information including gene names were obtained from the HUGO Gene Nomenclature Committee’s (HGNC) human genome database, comprising 42764 gene symbols and names, and 3245 gene families and sets as of the time of conducting this analysis. Information about the phenotypes, and the specific genes implicated in a syndrome was obtained from a survey of current literature in medical genetics and genomics, and from the Online Mendelian Inheritance in Man (OMIM) repository for authoritative information about human genes and genetic phenotypes.

### Algorithm 1

Pseudocode for a breadth-first algorithm for computing the set of chainlink genes that form various levels of voice chains {𝒱_*N α*_} for FOXP2.

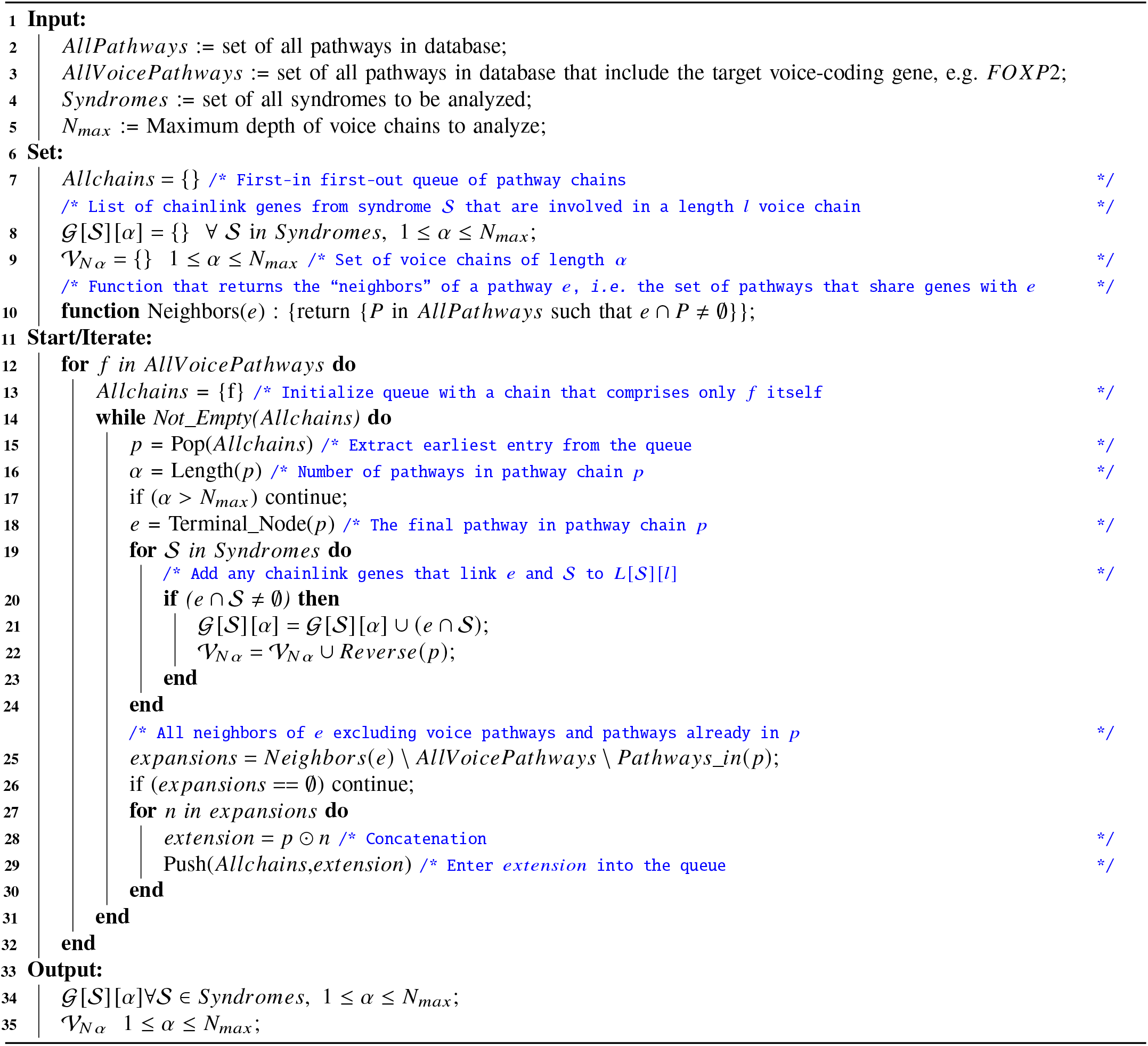

The FOXP2 gene chosen for this analysis has been strongly implicated in speech and language disorders [4], [5], including monogenic speech disorders. The cytogenetic location (chromosome locus) of this gene is 7q31.1. Mutations in this gene are known to cause speech and language disorder Type 1, also called “Autosomal dominant speech and language disorder with orofacial dyspraxia”. The phenotype description and known molecular basis for this disorder can be found under OMIM entry SPCH1:602081. The FOXP2 gene encodes for the protein “Forkhead Box Protein P2” [6]. This protein is a transcription factor – it controls the activity of other genes. It binds to the DNA of the genes that it controls through a region known as a Forkhead Domain. It thus plays a critical role in several protein-coding and other biological pathways, and has been well studied [7]. A more detailed summary of this gene can be obtained from the Human Protein Atlas [8].

The ensemble of pathways used for this analysis was obtained from the Carcinogenic Potency Database (CPDB), described on its website as “a single standardized resource of the results of 45 years of chronic, long-term carcinogenesis bioassays.” Its current database of human biological pathways contains 4319 pathways and their gene compositions. This database has been used extensively in the medical literature and was chosen in this case for illustrative purposes, since there is (importantly) no inherent bias towards the speech phenotype in it. In this database, there is only one pathway that contains the gene FOXP2. This is the Adenoid Cystic Carcinoma (ACC) pathway, which contains 63 genes, listed below for reference:

## Gene membership of the ACC pathway

AKT1 ARID1A ARID4B ARID5B ATM ATRX BCOR BCORL1 BRCA1 BRD1 CEBPA CMTR2 CNTN6 CREBBP CTBP1 DTX4 EP300 ERBB2 ERBIN FBXW7 FGF16 FGFR4 FOXO3 FOXP2 H1-4 H2AC16 HRAS IL17RD INSRR JMJD1C KANSL1 KAT6A KDM6A KDM6B KMT2C MAGI1 MAGI2 MAML3 MAP2K2 MAX MGA MORF4L1 MYB MYBL1 MYC MYCBP MYCN NCOR1 NFIB NOTCH1 NSD1 PIK3CA PRKDC PTEN RAF1 SETD2 SMARCA2 SMARCE1 SMC1A SRCAP TLK1 TP53 UHRF1

Table I documents the voice chains found for a set of 75 documented microdeletion syndromes. This range excludes chromosome 21, for which sufficient documentation was not found in the literature. The analysis was conducted for voice chains up to length 2 (Level-2). The number of pathways that a gene connects to (in general, inclusive of connections to the ACC pathway of FOXP2) is written as its subscript. The second column in Table I lists the corresponding speech phenotypes. References to the phenotypic description are cited. Where no reference is cited, the information reflects that found in the OMIM record for the syndrome. Where possible, OMIM references are used for brevity. The genes in the voice chains that have been implicated for the syndrome’s phenotypic expression in prior studies are shown in bold font.

**TABLE I:**
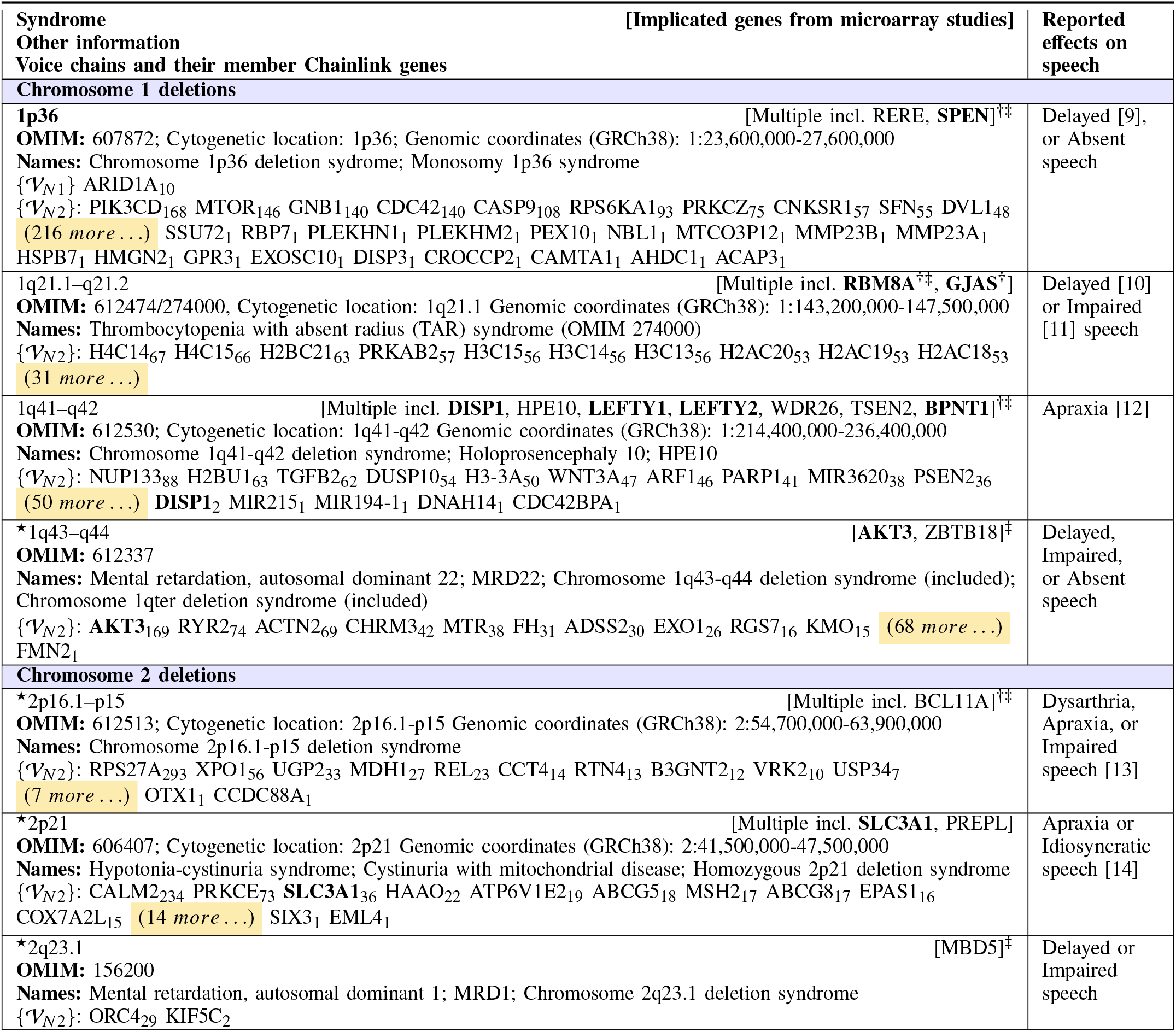
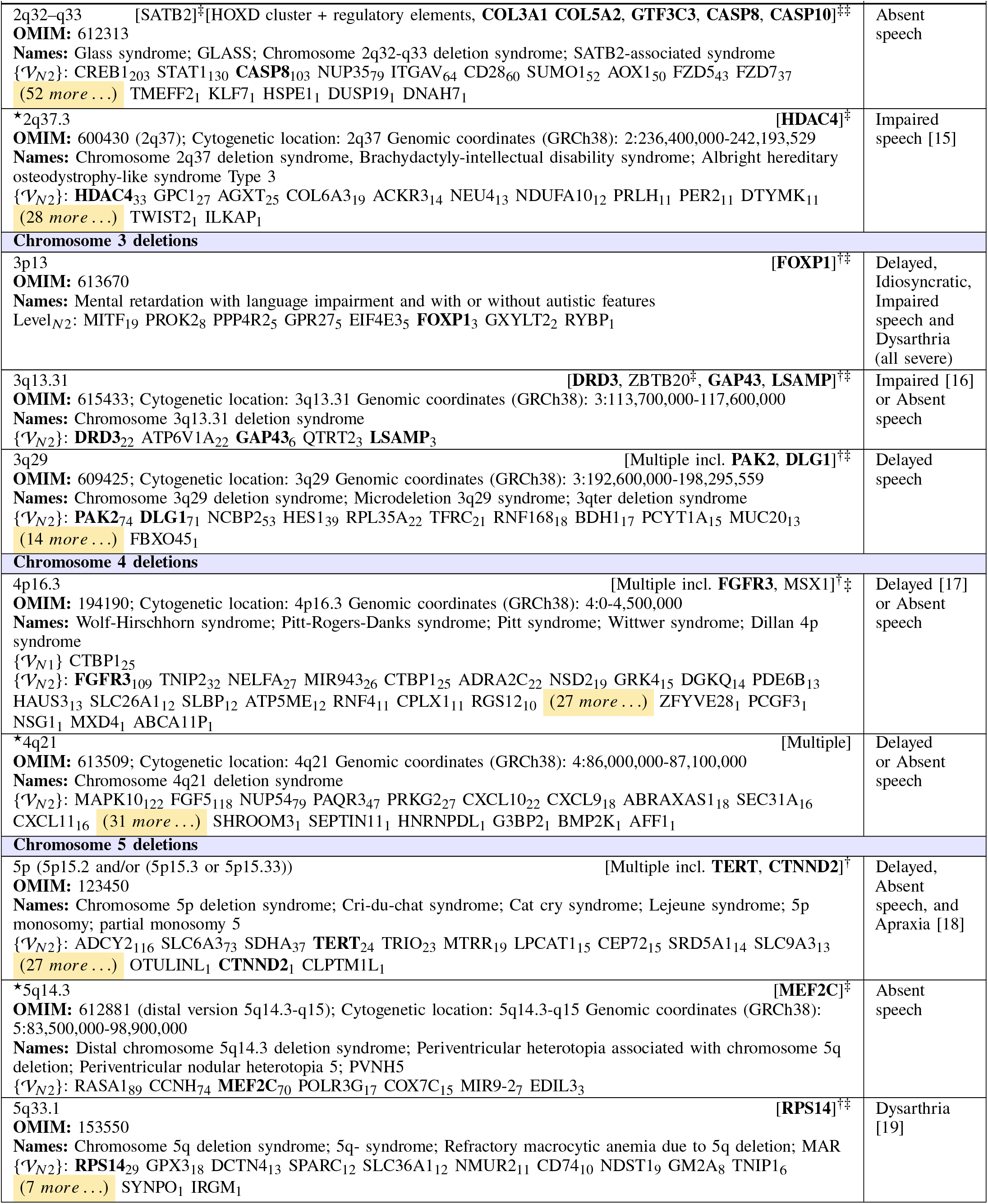
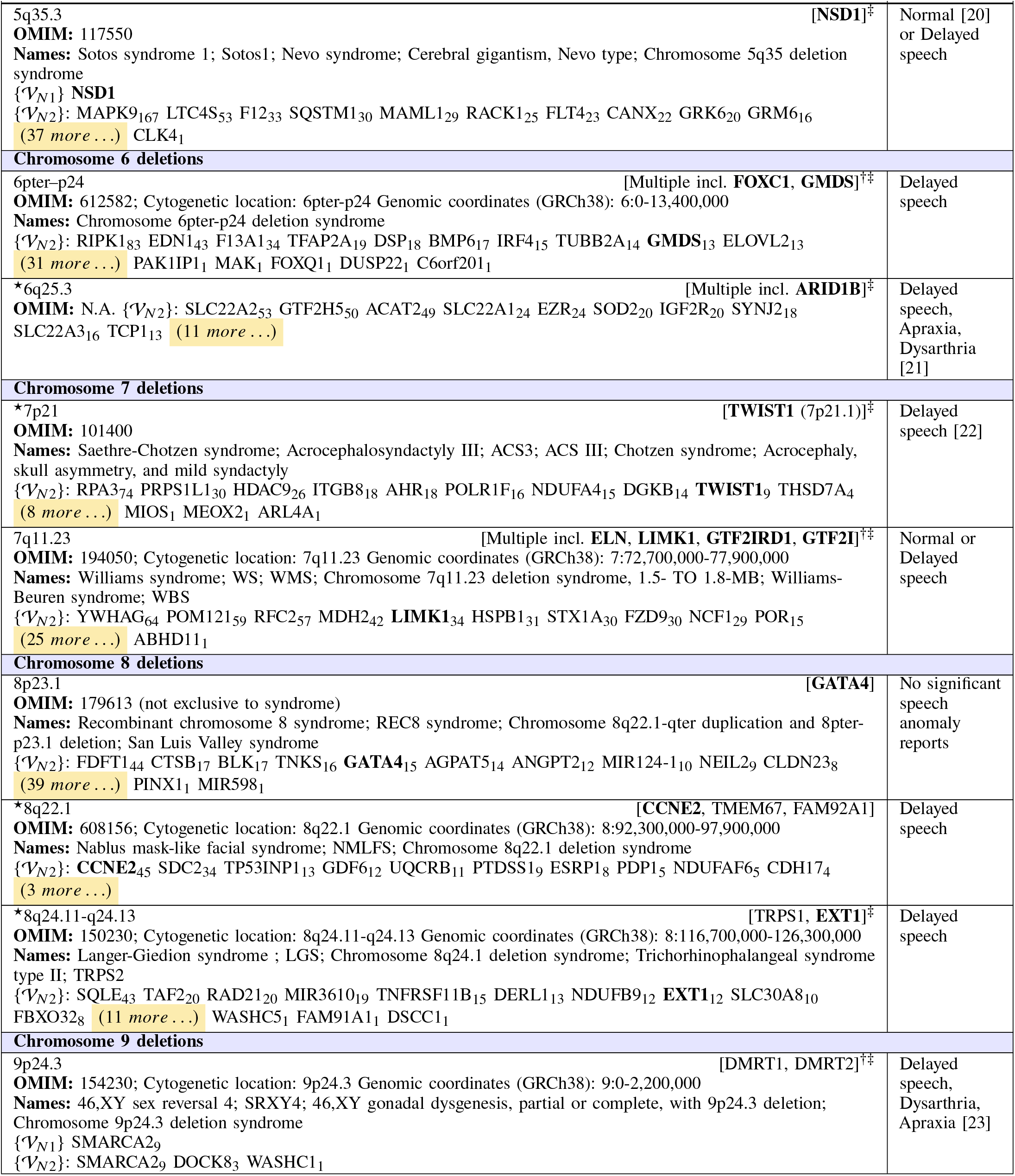
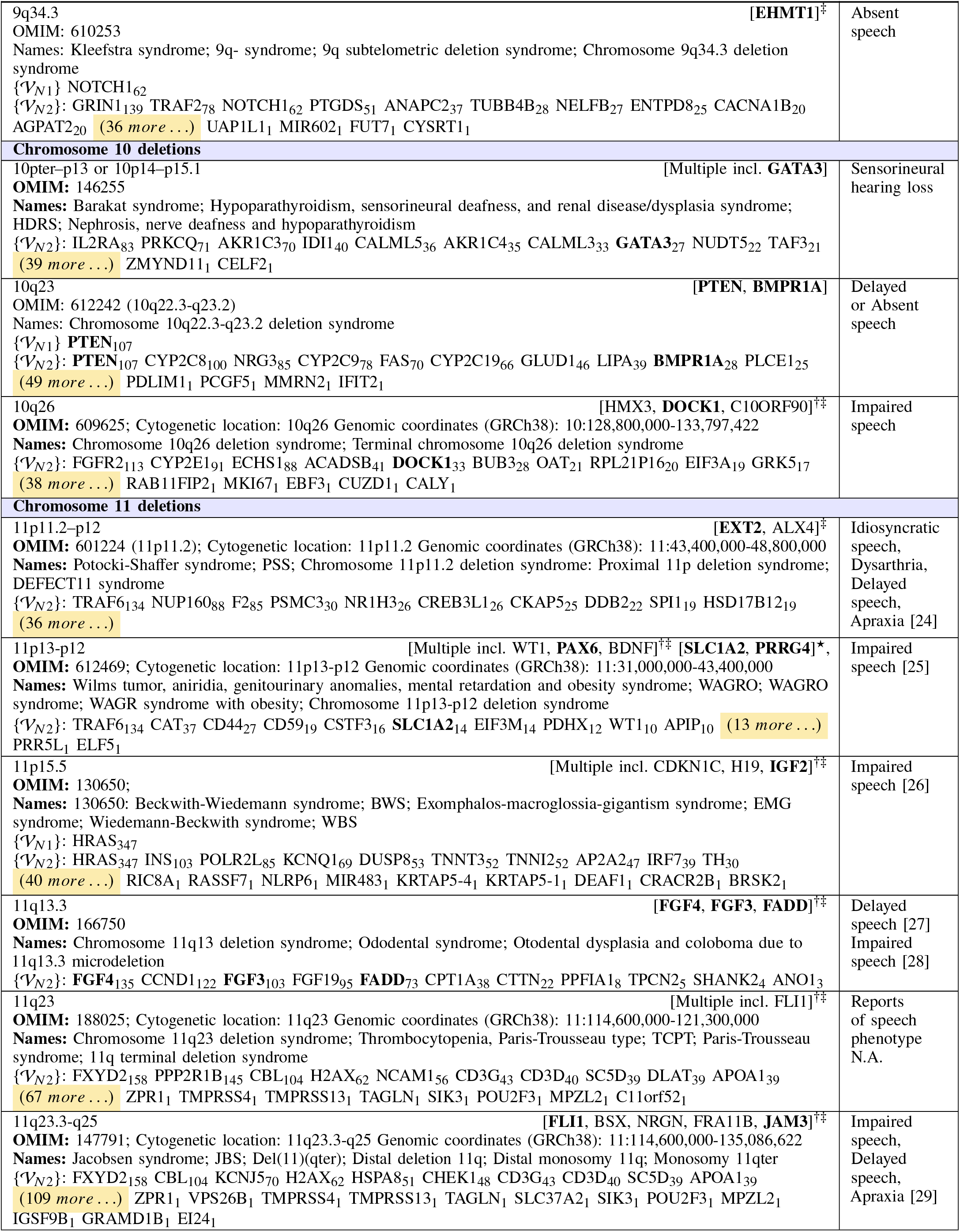
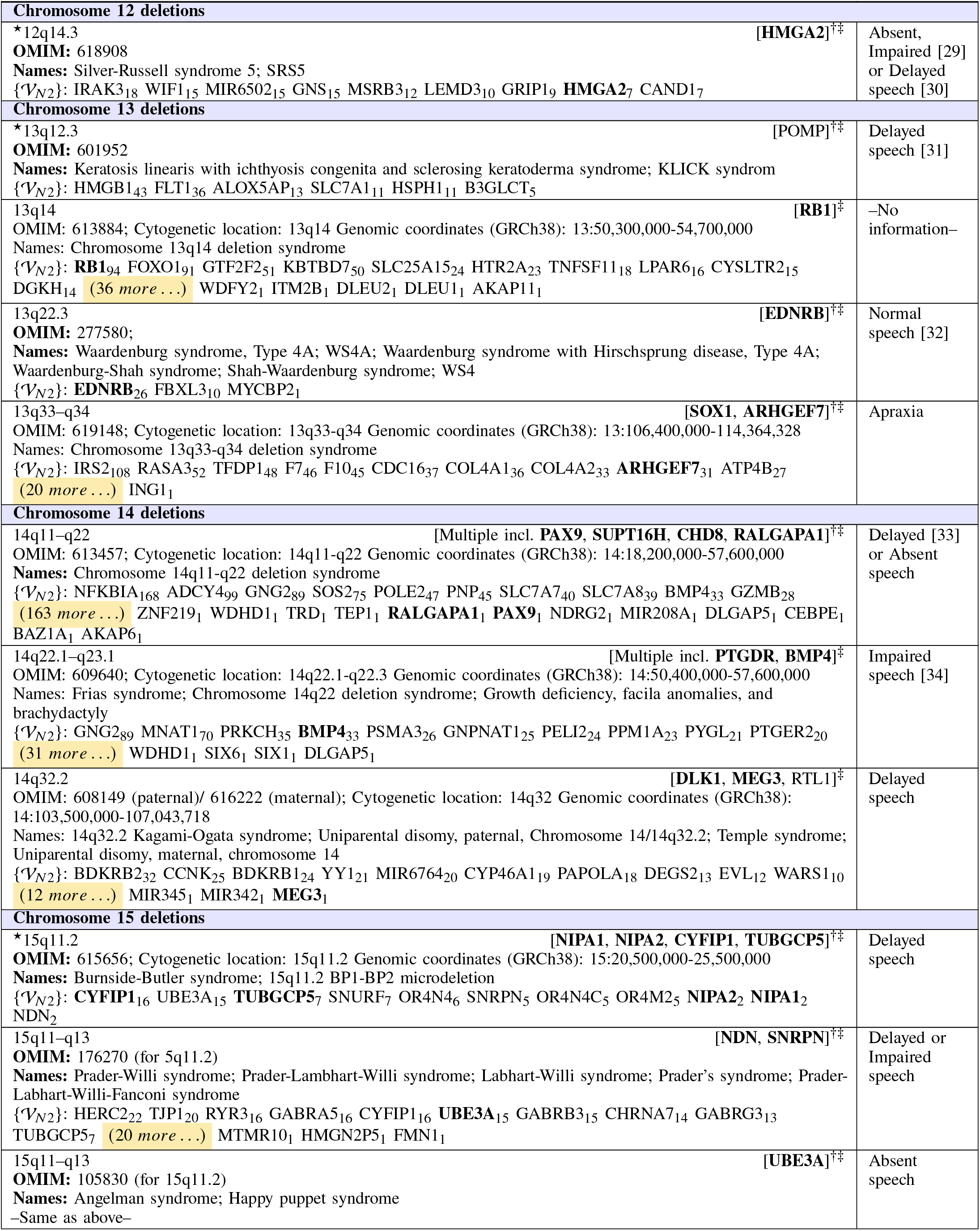
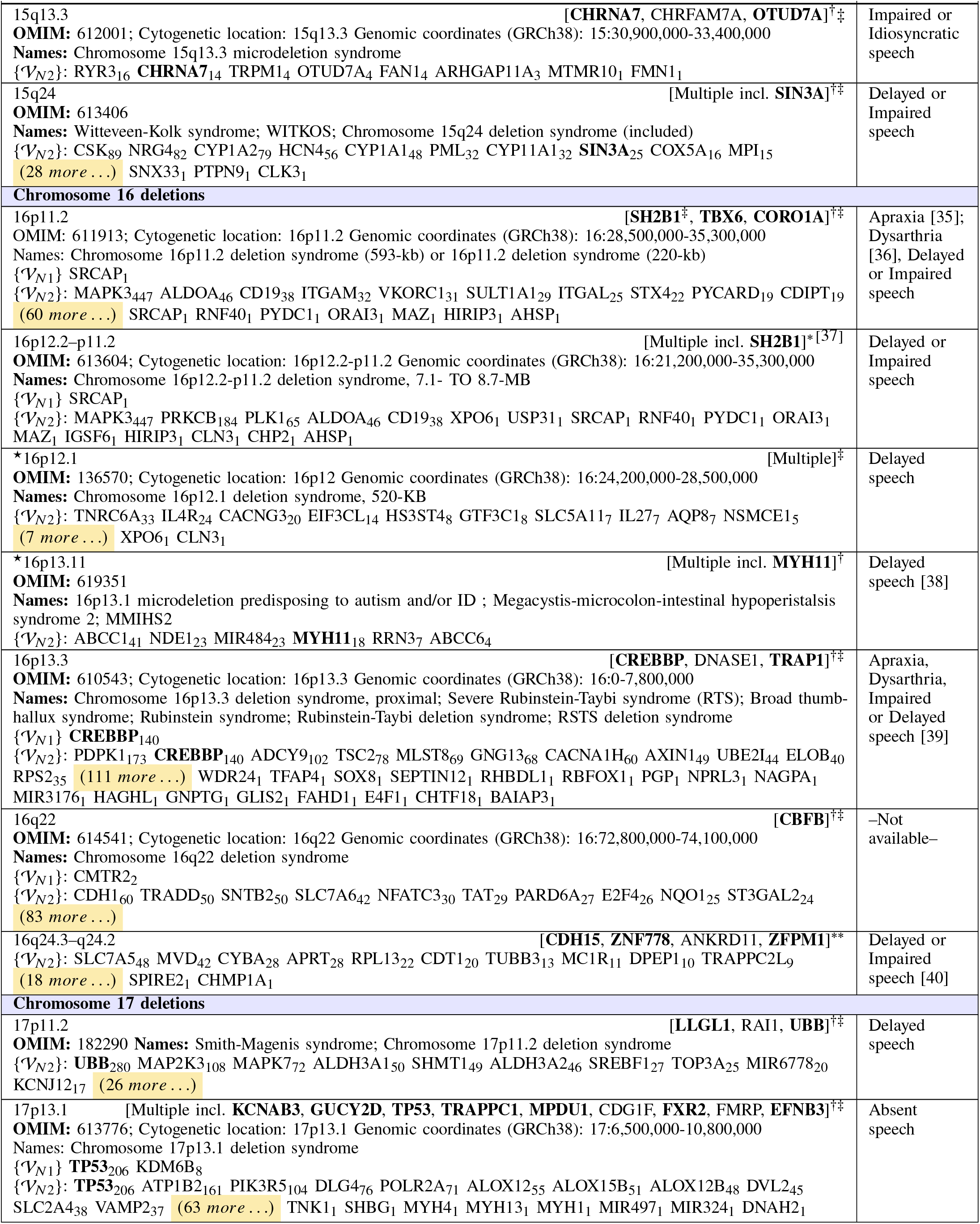
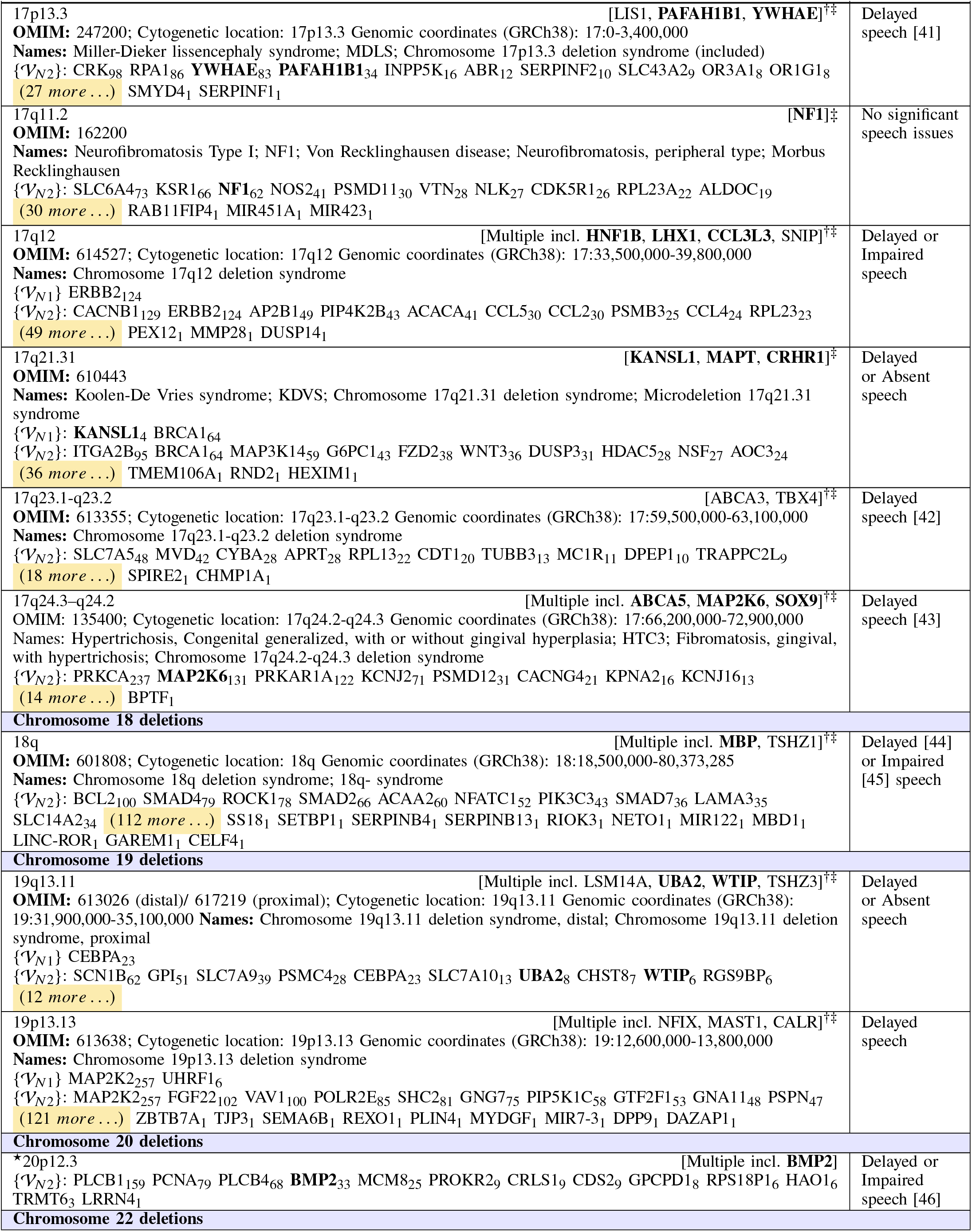
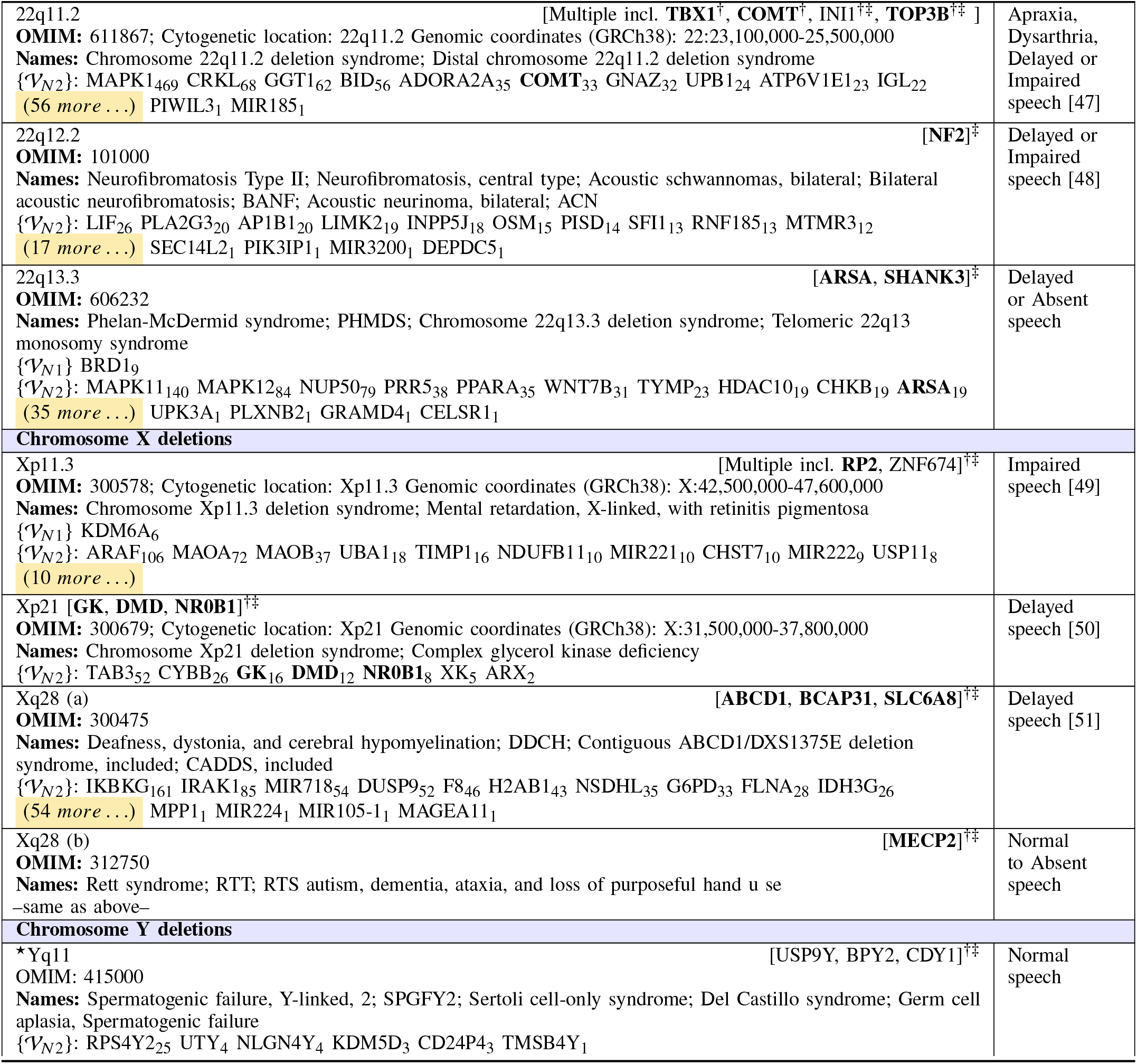
Chainlink genes for Level-1 and Level-2 voice chain ensembles for 75 chromosomal microdeletion syndromes. Each chain forms a link from the gene shown to the ACC biological pathway of FOXP2, and has been automatically derived. The total number of pathways that each gene influences independently is shown as a subscript to its name. Observed phenotypic effects on speech are given in the last column. When references are not cited, the information reflects that in the OMIM records for the syndrome. **Sources:** †: [52], ‡: [53], †‡: From OMIM records, ★: [25], ∗∗: [40]

## IV. Inferences

A wealth of conclusions can be drawn from Table I.

### 1) Voice chains as predictors of speech phenotype

Of the 75 syndromes analyzed, voice chains were found to exist for all. By our hypothesis, this would imply that in all cases, there is a potential for voice to be affected. In three of these (11q23, 13q114 and 16q22) no information about the speech phenotype was found. Only the remaining 72 syndromes are considered in the analysis below.

a. The incidence of speech pathologies (including *all* forms of pathologies) among the general population is reported to be about 5% [54], and between 2.3% and 24.6% among children [55]. Of the 72 syndromes, 18 had both level-1 and level-2 voice chains, while 54 had only level-2 chains. Occurrence of speech aberrations was reported for all 18 syndromes with level-1 chains, and for all but three (13q22.2, 17q11.2, and Yq11) of the 54 syndromes with only level-2 voice chains. Thus voice chains correlate highly with the existence of speech anomalies. In addition, we note that there were four syndromes (5q35.3, 7q11.23, 13q22.3 and X28(b)) for which studies have reported a range of phenotypes ranging from normal to completely absent speech. Only one of these syndromes (5q35.3) has a level-1 chain. However, given the inconclusiveness of the phenotypic reports, definitive conclusions cannot be made based on these syndromes. We consider these to be associated with aberrant speech for our analysis.
b. For each syndrome, studies so far have implicated some genes as being especially important, *i*.*e. implicated*. We find that of the 69 syndromes with reported speech anomalies, 2 do not have a specific implicated gene. These are the entries that simply say “Multiple” (4q21, 16p12.1). Of the 67 remaining that *do* have specific implicated genes, we note that in only 8 syndromes (2p16.1–p15, 2q23.1, 9p24.3, 11q23, 13q12.3, 17q23.1-q23.2, 19p13.13, Yq11), none of the implicated genes appear in the two levels of voice chains shown. Thus, in 59 of a total of 67 cases (where implicated genes are present), the implicated genes impact FOXP2 pathways, and the corresponding phenotypes show speech anomalies. However, in 2 of the three syndromes for which no aberrant speech has been reported, implicated genes are indeed found on the voice chains. The mere presence of an implicated gene on a voice chain may not be predictive of speech anomalies by itself. This may support the hypothesis that there may be factors other than the implicated gene that influence the phenotype for the syndrome.

### 2) Why are there no instances of missing voice chains?

Are voice chains redundant? The fact that there are no missing voice chains is linked in part to the size of the syndromic regions. Our database comprises 4319 unique pathways, 1205 of which are linked to the pathway that carries the FOXP2 gene (labelled 1422 in the ACC pathways database referenced earlier), and include 11746 genes. Thus, a randomly chosen gene from the *entire human genome* of 42.7k genes (as in the HGNC Human Genome database) has a 27.5% chance of being on a pathway that links to the 1422 pathway. The shortest microdeletion considered (2q23.1) includes 9 genes, and thus has a 94.44% probability of having a level-2 voice chain purely by chance. The second shortest pathway includes 26 genes and has a 99.98% probability of having a level-2 voice chain by chance. The remaining pathways are larger (in terms of the number of genes), and it is virtually impossible for them *not* to have a level-2 chain. As a result, it is realistic to expect that, as a consequence of the density with which the 1422 (FOXP2) pathway is linked to other pathways, *any* syndrome arising from genetic aberrations that includes even a moderately sized set of genes will have an effect on voice. More generally, any factor that influences gene function may be expected to ultimately affect voice.

The above argument assumes that the genes in a microdeletion region are randomly chosen. The mean of the fraction of genes in a microdeletion that appears in any voice chain is observed to be 28.79% with a variance of 0.014, indicating concordance with the assumption of randomness. A secondary implication is that the likelihood of adjacent genes in the same cytogenetic region to belong to a voice chain is independent of one another.

A second hypothesis is that the *strength* of the influence could potentially be related to the *level* of the voice chain – level-1 voice chains may lead to more severe effects on speech. This is examined below.

### 3) Level of voice chain and degree of influence

18 of the 72 syndromes considered have Level-1 voice chains, which means genes from these microdeletion regions also feature in the pathway on which FOXP2 also appears. With this, the speech phenotype is expected to be relatively more severe than for those syndromes for which Level-1 voice chains do not exist. Here we ignore the secondary effects of other phenotypes such as intellectual disability and craniofacial anomalies, a highly simplifying assumption. The syndromes with Level-1 voice chains, their corresponding genes and observed speech phenotypes are listed below:

|  |  |
| --- | --- |
| 1p36:ARID1A:Absent+Delayed | 4p16.3:CTBP1:Absent+Delayed |
| 5q35.3:NSD1:Delayed+Normal | 9p24.3:SMARCA2:Apraxia+Delayed+Dysarthria |
| 9q34.3:NOTCH1:Absent | 10q23:PTEN:Absent+Delayed |
| 11p15.5:HRAS:Impaired | 16p11.2:SRCAP:Apraxia+Delayed+Dysarthria+Impaired |
| 16p12.2–16p11.2:SRCAP:Delayed+Impaired | 16p13.3:CREBBP:Apraxia+Delayed+Dysarthria+Impaired |
| 16q22:CMTR2:— | 17p13.1:KDM6B,TP53:Absent |
| 17q12:ERBB2:Delayed+Impaired | 17q21.31:BRCA1,KANSL1:Absent+Delayed |
| 19q13.11:CEBPA:Absent+Delayed | 19p13.13:MAP2K2,UHRF1:Delayed |
| 22q13.3:BRD1:Absent+Delayed | Xp11.3:KDM6A:Impaired |

If we rank order the phenotypes by occurrence count, we get the following order: Normal(1) < Dysarthria(3) = Apraxia(3) < Impaired(6) < Absent(8) < Delayed(13)

We see that together, the absence of speech, delay in speech development, and impairment of speech – all relatively stronger phenotypes – constitute the dominant set. This is consistent with our expectation that shorter voice chains to FOXP2 may have more intense effects.

Analysis of level-2 chains is more complicated. While a syndrome can only have one level-1 voice chain, it can potentially have a large number of level-2 chains. Consequently, the number of chainlink genes may be expected to be much larger for level-2 chains than for level-1 chains. The average number of chainlink genes for level-1 pathways is only 1.166, whereas that number for level-2 pathways is 45.04. As a result, although the increased length of the level-2 chains may be expected to reduce the direct influence of any gene, both, the increased number of genes and the increased number of level-2 paths through which any individual gene may connect to FOXP2 can compound this influence.

To evaluate this hypothesis, we assigned a 0-4 rating to the speech phenotypes by increasing degree of severity as follows: Normal:0, Dysarthria:1, Apraxia:2, Idiosyncratic:2, Impaired:2, Delayed:3 and Absent:4.

Each syndrome was assigned a speech score, comprising an average of the ratings of the phenotypes reported for a syndrome. We note that all 18 syndromes with level-1 chains also included level-2 chains. The remaining 54 (of 72) syndromes only had level-2 chains.

The average phenotype score for syndromes with both level-1 and level-2 chains was found to be 2.85 with a variance of 0.64. The average score for syndromes with only level-2 chains was found to be 2.40 with a variance of 0.75. A t-test shows the difference to be significant with *p* < 0.05, showing that in spite of the increased connectivity, level-2 chains have significantly lower influence than level-1 chains. Nonetheless, the increased variance of phenotypes for level-2 chains can result in serious impairment phenotypes. For instance, syndromes 14q11-q22, with 185 genes in level-2 chains and 4q21 with 47 genes in level-2 chains both are associated with absent speech. On the other hand, all of the syndromes that have no reported influence on speech also only have level-2 chains, but no level-1 chains.

However, even such an analysis would be naїve, since they do not consider the secondary effects of other phenotypes, such as intellectual disability, craniofacial anomalies, hearing disabilities, psychomotor retardation (all of which affect speech), which must be taken into account. Our current analysis is agnostic of the co-occurrence and secondary effects of other factors, which are likely to make the prediction of voice anomalies more certain, and the predicted effects stronger.

### 4) Identifying candidates genes for further investigation

Using only level-1 chains as illustrative examples, we see that voice chains can be useful in identifying candidate genes for further investigation in the context of speech phenotypes. As evidence of this, we consider the testimony of the genes that have been implicated for the syndromes through independent studies.

- 1p36 (Likely candidate (lc): ARID1A): Although not implicated for this syndrome in studies so far, ARID1A is located in 1p36.11, a region frequently deleted in human cancers [56]. Disruption in its function may lead to co-occurrence of oncological and speech phenotypes. This hypothesis is verifiable.
- 5q35.3 (lc:**NSD1**) The gene NSD1 appears in the Level-1 chain and is also an implicated gene. Ideally, this should not be a candidate for further investigation. However, paradoxically, while effects on speech are expected, the literature reports normal speech for some subjects for this phenotype. The reason for this is not clear yet and potentially warrants investigation.
- 11p15.5 (lc:HRAS) Although not implicated, and although two studies cited under OMIM: 130650 for this syndrome explicitly mention HRAS as *not* significant, HRAS has nevertheless been independently found to be extremely significant in RASopathy and cancer studies, e.g. [57]. Its role in this syndrome needs to be re-evaluated, given its influence on 347 biological pathways and the strong speech phenotype.
- 16p11.2 and 16p12.2–16p11.2 (lc:SRCAP) Although not implicated, it connects to only one other pathway in the ensemble, and that is the ACC pathway of FOXP2. The effects on speech are expected to be strong if this gene is aberrant. This gene may be implicated in further investigations.
- 17p13.1 (lc:KDM6B) Speech is absent in this syndrome. The gene TP53 is implicated which also appears at Level-1 is associated with 206 pathways. KDM6B is the only other gene in the level-1 voice chains, and connects to only 8 other pathways. It is likely that this gene also plays a strong role in the speech phenotype and merits investigation.
- 17q12 (lc:ERBB2) The gene ERBB2 is associated with 124 pathways. It is a well-known oncogene [58], in that perturbations in its function have been observed to have deleterious effects. If it is also connected to FOXP2, then its appearance in the voice chain allows a surprising hypothesis – that biomarkers of some oncological conditions may also be present in voice.
- 19p13.3 (lc:MAP2K2,UHRF1) MAP2K2 and URHF1 are not implicated. However their presence in the level-1 chain warrants investigation, especially for MAP2K2 which influences 257 pathways. Prompted by this, a literature search reveals that MAP2K2 *has* been implicated in this syndrome recently [59], although this is not on the OMIM records, which were largely consulted for this study.
- 22q13.3 (lc:BRD1) The gene BRD1 is not implicated, and appears in 9 pathways only, but the speech phenotype is severe in this syndrome. This warrants the investigation of BRD1 independently in speech phenotypes. A literature search reveals that BRD1 is indeed strongly associated with brain development and susceptibility to both schizophrenia and bipolar affective disorder [60], and consequent effects on speech are highly likely.
- Xp11.22 (lc:SMC1A) Although SMC1A is not implicated, It appears in 33 pathways. The speech phenotypes are severe and the gene warrants investigation for this phenotype. A recent report in the literature has implicated it in severe intellectual disability and therapy-resistant epilepsy in females [61]. The former is known to be associated with severe speech anomalies.
- Xp11.3 (lc:KDM6A) Although not implicated, KDM6A warrants investigation. In the literature, it is independently known to be associated with delayed speech, and psychomotor development [62].

### 5) Expression of the speech phenotype

The observation that deletions of genes on *all chromosomes* ultimately results in the expression of speech anomalies carries significance. From a much broader perspective, this overwhelmingly suggests that the speech phenotype may be an emergent capability, regulated and supported by the action of multiple concurrent biological pathways. There may be no single gene or genes (on select chromosomes) that may code for speech capabilities per-se, and FOXP2 may be one of a few genes that may *consolidate and regulate* the speech and language related emergent effects. It may be that genes directly code for structural elements in the range of phenotypes, while other properties such as speech and language abilities, are emergent from the co-ordination of these (and epigenetic) factors.

A more prosaic argument in favor of a phenotype being an emergent effect can also be presented. Within the ensemble of syndromes analyzed, there are three kinds of of cause-and-effect relationships: a) syndromes with *physical* structures of the vocal tract (e.g. craniofacial anomalies that include cleft palate, changes in lip shape etc.), which adversely affect the biomechanical aspects of voice and speech production, b) syndromes in which auditory and motor functions are compromised, and c) syndromes that affect the normal functions of the brain, causing cognitive, learning, memory, and other issues that are in turn likely to lead to speech problems. In no case do we see only speech aberrations in isolation of these phenotypes. The associations between speech and other phenotypes have in fact been ubiquitously observed, e.g. [38]. This supports the categorization of speech abilities as an emergent property of an ensemble of factors (including other phenotypes), rather than a primary phenotype that is likely to be expressed directly by a gene.

The emergence hypothesis is further supported by the fact that both, monosomies *and* trisomies in the case of autosomal aneuploidies have deleterious effects. This suggests the existence of an unstable equilibrium state where “normalcy” of phenotype is emergent from the correct functioning of all pathways, and disrupted by a disruption in any. Perturbation of this state from equilibrium in any sense seems to cause not only anomalies, but almost exclusively deleterious ones.

The implications of an emergence genesis for some phenotypes are important to consider: since emergent properties are not as tightly bound to the direct expression of genes, it may be possible to restore aberrant functions in these by adjustment of the overall properties of the pathway ensemble involved. In simpler words, the “cure” for a genetic condition may not necessarily only lie in restoring gene function, but may possibly also be in restoring some quantifiable equilibrium.

## V. Conclusions

The hypothesis that the existence of voice chains is correlated with speech phenotypes is adequately validated by the statistical analysis presented in this paper. We also illustrate how the methodology presented can potentially provide leads to specific genes that might be candidates for further investigation in the context of speech phenotypes and microdeletion syndromes. The methodology itself may potentially be generalized and extended to reveal the potential effects of other diseases with genetic basis, and of other factors that influence gene function in some manner, on speech.

## VI. Acknowledgment

This material is based upon work supported by the U.S. Army Research Office and the U.S. Army Futures Command under Contract No. W911NF-20-D-0002. Its content does not reflect the position or the policy of the U.S. Army and no official endorsement should be inferred.

